# Deciphering the Effect of Melittin on Murine Cervical Cancer Cells Based on Transcriptomic Investigation

**DOI:** 10.64898/2026.07.29.741659

**Authors:** Jiang Jianrong, Zhang Yuwei, Wang Mengyi, Zhang Ronghua, Li Yueliang, Qiu Jianfeng, Chen Dafu, Yan Tizhen, Guo Rui, Liu Yanhui

## Abstract

Melittin, a highly active natural antimicrobial peptide derived from honeybee venom, holds immense pharmacological potential against solid tumors. However, its specific anti-tumor efficacy and transcriptomic dynamics in cervical carcinoma remain to be systematically characterized. This study evaluated the anti-cancer properties of melittin on murine U14 cervical cancer cells following by transcriptomic investigation of the underlying mechanism. Phenotypic evaluations revealed that melittin potently inhibited U14 cell viability, while wound healing assays demonstrated a profound, dose-dependent suppression of cellular migration, culminating in near-complete migratory arrest at high concentrations. Furthermore, flow cytometry quantified a dramatic, dose-dependent surge predominantly in late apoptotic cell populations. These apoptotic events were structurally corroborated by scanning electron microscopy (SEM), which revealed severe plasma membrane perforation and morphological exhaustion. Enrichment analyses indicated that the physical membrane disruption inflicted by melittin translated into a severe metabolic crisis, marked by a global suppression of ribosomal biogenesis and mitochondrial oxidative phosphorylation. Furthermore, melittin profoundly repressed the Tie2-mediated angiogenic pathway (*Etv2* down-regulation) while triggering the lethal hyperactivation of the AP-1 transcriptional stress complex (*Jun*, *Fos*, *Fosb*). Collectively, these findings elucidate the pharmacological networks underlying melittin’s cytotoxicity, providing solid molecular evidence for its development as a natural therapeutic agent against cervical cancer.

## Introduction

Cervical cancer is a malignant tumor originating from the epithelial cells of the cervix ^[1]^. The cervix constitutes the lower portion of the uterus, connecting it to the vagina. Tumors form when cervical cells undergo abnormal DNA alterations and proliferate uncontrollably ^[2]^. Globally, there are approximately 661,000 new cases of cervical cancer and 348,000 related deaths annually. In China, the incidence and mortality rates of cervical cancer rank among the highest worldwide, and the disease presents a trend toward younger demographics, posing a severe threat to women’s health and lives ^[3]^. Although the widespread administration of human papillomavirus (HPV) vaccines and advancements in screening technologies have significantly reduced the incidence of cervical cancer, it remains a leading cause of cancer-related mortality among women in developing countries due to limited medical resources and inadequate screening coverage ^[4]^. Consequently, cervical cancer continues to be a pressing global public health challenge. Currently, the primary treatment modalities for early-stage cervical cancer include surgical resection, radiotherapy, and chemotherapy, whereas concurrent chemoradiotherapy or targeted therapy is employed for advanced or recurrent cases ^[5]^. However, traditional chemotherapeutic agents, such as cisplatin and paclitaxel, are associated with severe toxic side effects. Furthermore, tumor cells are prone to developing multidrug resistance, giving rise to treatment failure and disease progression ^[6–7]^. In recent years, despite the promise shown by immune checkpoint inhibitors (e.g., protein 1/programmed death-ligand 1 (PD-1/PD-L1) antibodies) for certain patients, the overall response rate remains below 20%, and the treatment costs are prohibitively high ^[2,8]^. Hence, developing novel anti-cervical cancer drugs with high efficacy and low toxicity, as well as deeply elucidating their mechanisms of action, is of significant clinical and scientific value.

Melittin, constituting approximately 50% of the dry weight of bee venom, is its principal active component. As a cationic amphiphilic peptide composed of 26 amino acids, melittin is considered a highly potent candidate for cancer therapy due to its robust antitumor activity ^[9–11]^. Its antitumor mechanism relies not only on its membranolytic properties but also on its ability to effectively eliminate tumor cells by inducing necrosis and apoptosis ^[12,13]^. To date, certain progress has been made in research regarding the anti-cervical cancer effects of melittin. Studies have revealed that in cervical cancer models, melittin exhibits significant dose-dependent cytotoxicity, inducing G_0_/G_1_ cell cycle arrest and mitochondria-mediated apoptosis ^[14]^. Furthermore, melittin can specifically downregulate the expression of the E6 and E7 oncoproteins of high-risk HPV types, such as papillomavirus 16 and human papillomavirus 18 (HPV16 and HPV18), thereby selectively killing HPV-positive cervical cancer cells ^[15]^. Nevertheless, current research on the efficacy of melittin against cervical cancer remains in its preliminary stages, and the underlying molecular regulatory networks require further in-depth elucidation.

In the present work, using the murine cervical cancer U14 cell line as a model, we integrated macroscopic phenotypic characterizations, including cell viability, flow cytometry-based apoptosis assays, wound healing, and ultrastructural evaluations via scanning electron microscopy (SEM) with global transcriptome sequencing (RNA-seq) to systematically screen for differentially expressed genes (DEGs) following melittin treatment. Rather than focusing solely on isolated metabolic pathways, our multi-level bioinformatic evaluation, utilizing Gene Ontology (GO), the Kyoto Encyclopedia of Genes and Genomes (KEGG), Reactome, Gene Set Enrichment Analysis (GSEA), and protein-protein interaction (PPI) network topology maps a comprehensive, system-wide functional shift. Specifically, our primary focus highlights melittin ’ s capacity to couple direct physical membrane disruption with a complete paralysis of ribosomal translation and oxidative phosphorylation, the blockade of TEK receptor tyrosine kinase (Tie2)-mediated angiogenic migration, and the lethal hyperactivation of the activator protein 1 (AP-1) transcriptional stress interactome. This study aims to thoroughly elucidate the multi-targeted molecular mechanisms of melittin against cervical cancer at both the structural and transcriptional levels, thereby providing a solid theoretical foundation and robust experimental support for its clinical translation and application.

## 1 Materials and Methods

### 1.1 Cell Culture and Melittin

Murine cervical cancer cells (Xiamen Immocell Biotechnology), maintained at the Laboratory of Host-Pathogen Interaction Mechanisms and Precision Medicine, College of Bee Science of Fujian Agriculture and Forestry University, were cultured in complete medium (Thermo Fisher Scientific, Waltham, MA, USA) at 37 ℃ in a 5% CO_2_ atmosphere. Melittin (purity ≥ 95%) was purchased from Selleck (USA).

### 1.2 Cell Viability and Colony Formation Assays, and Dose Selection

Following the instructions, cell proliferation and viability were assessed using the Cell Counting Kit-8 (CCK-8) assay (Beyotime, Shanghai, China). Briefly, U14 cells were seeded into 96-well plates, incubated overnight, and treated with varying concentrations of melittin (0, 2, 4, 6, and 8 μg/mL) for 20 min. OD was measured at 450 nm to determine the IC50. For the colony formation assay, cells were seeded into 6-well plates 500 cells/well, exposed to the same melittin gradient for 20 min, and cultured in drug-free medium for an additional 7 days. Macroscopic colonies were fixed with 4% paraformaldehyde and stained with 0.1% crystal violet. On basis of the fact that melittin drastically suppressed the colony-forming capacity of U14 cells in a dose-dependent manner, alongside the CCK-8 viability data, 4 μg/mL was rigorously selected as the optimal intervention concentration for all subsequent transcriptomic and mechanistic analyses.

### 1.3 Wound Healing Assay

U14 cells were seeded into six-well plates and grown to full confluence. An artificial wound was created by scratching the cell monolayer linearly with a sterile 200 μL pipette tip. After being washed with PBS to remove cellular debris, the cells were cultured in medium containing melittin (0, 2, 4, 6, and 8 μg/mL). Images of the identical wounded areas were captured at 0 h and 16 h post-wounding using inverted phase-contrast microscope (Sunny Optical, China).

### 1.4 Annexin V-FITC/PI Apoptosis Assay

U14 cells treated with varying concentrations of melittin (0, 2, 4, 6, and 8 μ g/mL) were harvested and washed with cold PBS. The collected cells were resuspended in 1× binding buffer and double-stained with Annexin V-FITC and Propidium Iodide (PI) (Yeasen, Shanghai, China) in the dark according to the protocol. The stained cells were immediately acquired utilizing a flow cytometer, and the proportions of apoptotic cells across different sub-populations were quantified.

### 1.4 Transcriptome data source

In a previous study, melittin- and un-tretaed U14 cell samples were prepared and subjected to RNA isolation, cDNA library construction, and deep sequencing, followed strict quality control of raw reads, obtaining high-quality clean reads (https://doi.org/10.64898/2026.07.25.739748).

### 1.5 SEM detection

U14 cells treated with 4 μg/mL melittin or complete medium for 20 min were evaluated for gross morphological alterations utilizing an inverted microscope. For ultrastructural imaging, cells were fixed in 2.5% glutaraldehyde at 4°C overnight, post-fixed with 1% OsO_4_ for 1 h, and dehydrated through a graded ethanol series. Following critical-point drying and gold sputter-coating, the cell surface topography was visualized using a scanning electron microscope (Carl Zeiss, Oberkochen, Germany).

### 1.6 DEG screening and enrichment Analysis

DEGs between the un- and melittin-treated groups were screened using the DESeq2 package (v1.22.2), based on the cutoff criteria of an absolute log_2_ fold change (|log_2_FC|) ≥ 1 and *Q* value < 0.05. To decipher the biological implications of these transcriptomic alterations, GO enrichment was performed via the clusterProfiler package using a hypergeometric test.

To further map these DEGs into systemic biological pathways, we queried the KEGG (https://www.kegg.jp/) and Reactome (https://reactome.org/) databases, defining significance as *P*<0.05. Instead of relying solely on predefined DEGs, we also assessed global transcriptomic shifts using GSEA, with significant sets defined by an absolute normalized enrichment score (|NES|) > 1, nominal *P* value < 0.05, and false discovery rate (FDR) < 0.25. Finally, to identify core regulatory hubs within the altered pathways, a protein-protein interaction (PPI) network was constructed utilizing STRING (v11.5) and visualized in Cytoscape (v3.9.1).

### 1.7 qRT-PCR Analysis

Total RNA was isolated from the cervical cancer cells utilizing the Accurate RNA Extraction Kit (Accurate Biotechnology). Following the assessment of RNA yield and integrity via NanoDrop spectrophotometry and agarose gel electrophoresis, equal amounts of purified RNA were subjected to reverse transcription using the Hifair® III cDNA Synthesis Kit (Yeasen, Shanghai, China). To evaluate the transcript abundance of 6 selected genes (*Fos*, *Fosb*, *Jun*, *Nr1i2*, *Prdm1*, and *Egr1*), quantitative PCR (qPCR) was executed on a QuantStudio 3 system (Applied Biosystems) using SYBR Green chemistry (Yeasen, Shanghai, China). The amplification was performed in a 20 μL reaction volume with an initial denaturation at 95°C for 3 min, followed by 40 cycles of 95°C for 15 s, 58°C for 30 s, and 72°C for 15 s. All gene-specific primers (Table 1) were synthesized commercially (Sangon Biotech), with *Gapdh* employed as the endogenous reference. All assays were conducted in technical triplicates.

**Table 1.**
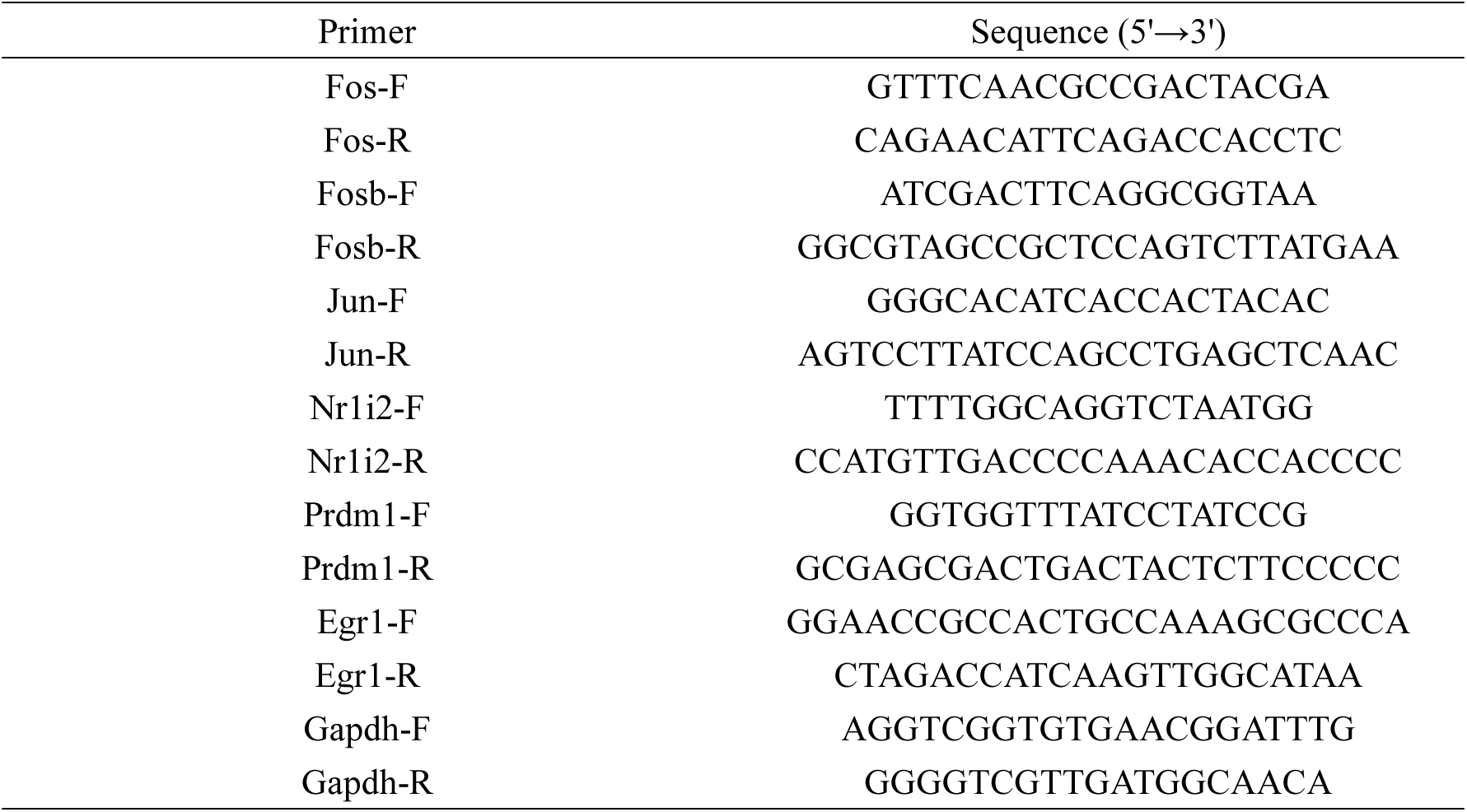
Sequences of Primers used in this study.

### 1.8 Statistical Analysis

Relative gene expression levels were quantified employing the2 ^−△ △ Ct^ method ^[16]^. All statistical analyses and graphical representations were executed utilizing GraphPad Prism 8 software. Quantitative data are presented as the Mean ± Standard deviation (SD). Statistical significance between the experimental and control groups was evaluated via Student’s *t*-test and Two-way ANOVA . Statistical significance was indicated as follows: ns, *P* > 0.05; *, *P* < 0.05; **, *P* < 0.01; ***, *P* < 0.001; and ****, *P* < 0.0001.

## 2 Results

### 2.1 Assessment of cell viability and clonogenic survival following melittin intervention

To characterize the cytotoxic effect of melittin, we first evaluated U14 cell viability using the CCK-8 assay. Quantitative absorbance (OD 450) measurements via a microplate reader revealed a dose-dependent reduction in viability following melittin exposure. Notably, while the 2 μ g/mL melittin treatment showed no significant inhibitory effect compared to the control, concentrations at or above 4 μg/mL induced a statistically significant decrease in cell viability (**Figure 1A**), from which the IC50 value was calculated (**Figure 1B**).

**Figure 1.**
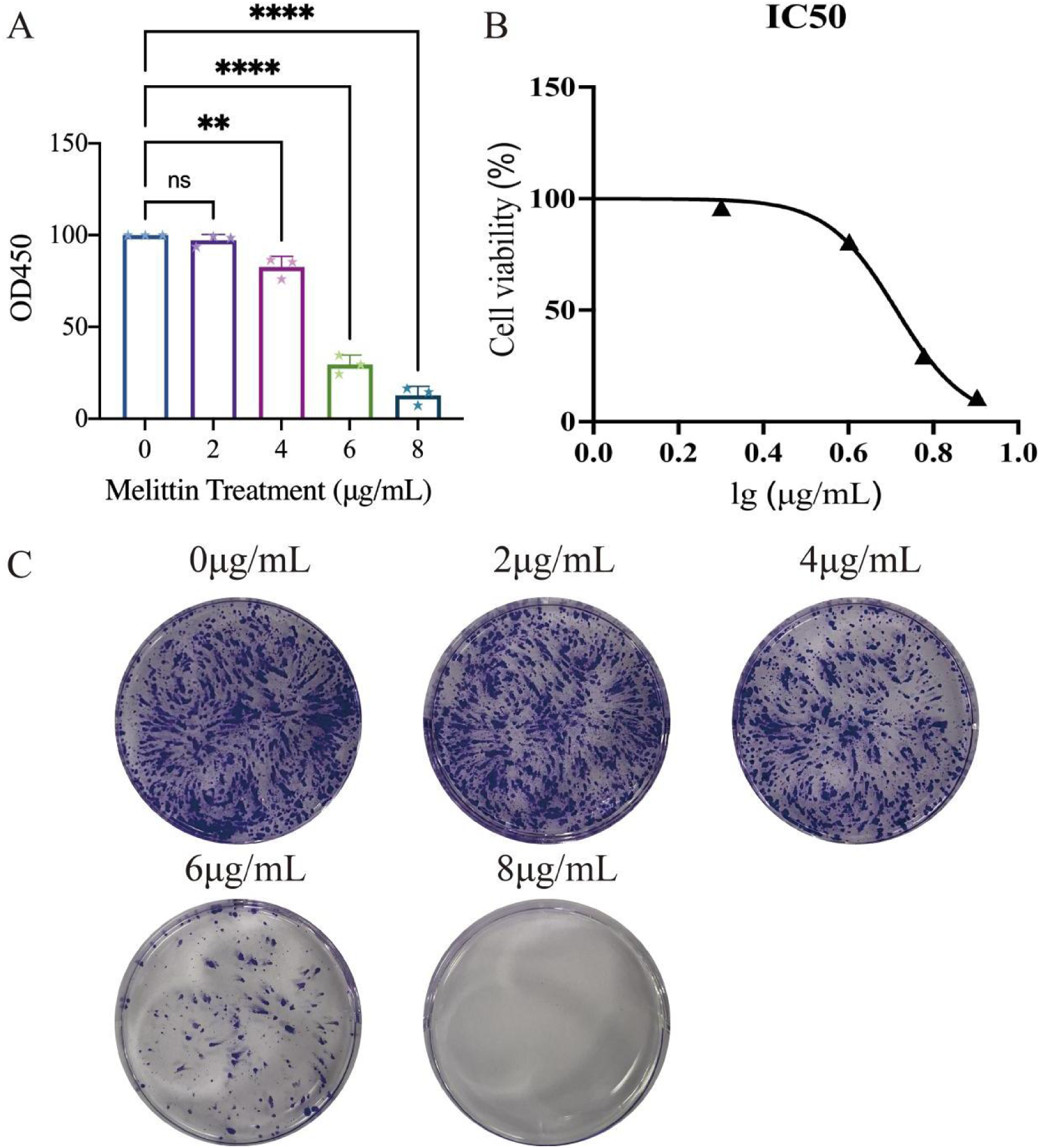
Melittin suppresses the short-term viability and long-term clonogenic survival of U14 cervical cancer cells. (A) Viability of cells treated with the indicated concentrations of melittin (0, 2, 4, 6, and 8μg/mL) assessed via CCK-8 assay. (B) Dose-response curve for determining the half-maximal inhibitory concentration (IC50) of melittin. (C) The colony formation assay. Data are presented as mean ± SD. ns, not significant; *P* < 0.01, **** *P* < 0.0001 compared to the un-treated control group (0μg/mL) (Student’s *t*-test).

Given that acute cytotoxicity assays only define immediate lethality, a colony formation assay was performed to ascertain the impact of melittin on the long-term proliferative potential of U14 cells. Melittin drastically suppressed the clonogenic capacity in a dose-dependent manner (**Figure 1C**). Control cultures (0 μg/mL) proliferated robustly, forming dense and macroscopic colonies. While the 2 μg/mL treatment largely retained colony formation, exposure to 4 μg/mL noticeably impaired the clonogenic capacity. This inhibitory effect progressed severely at 6 μ g/mL, resulting in drastically reduced colony numbers and sizes. Strikingly, intervention at 8 μ g/mL led to the complete eradication of clonogenic potential. Collectively, these phenotypic data robustly substantiate the potent anti-cervical cancer activity of melittin. Based on these viability and clonogenic survival analyses, 4 μg/mL was rigorously selected as the optimal intervention concentration for subsequent global transcriptomic investigations.

### 2.2 Assessment of cell migration following melittin intervention

To evaluate the effect of melittin on U14 cell migration, a wound scratch assay was performed. As depicted in Figure 4, the control group exhibited substantial cellular migration into the denuded zone, leading to significant wound closure after 16 hours. In contrast, melittin intervention markedly impeded the migratory capacity of U14 cells in a dose-dependent manner. While the 2 μg/mL treatment group showed partial wound healing, exposure to higher concentrations (4 and 6 μg/mL) severely restricted cell motility, resulting in noticeably wider remaining scratch areas. Strikingly, at the highest concentration of 8 μ g/mL, the scratch width remained virtually unchanged at 16 hours compared to the initial 0 h time point, indicating a near-complete arrest of cell migration(**Figure 2**).

**Figure 2.**
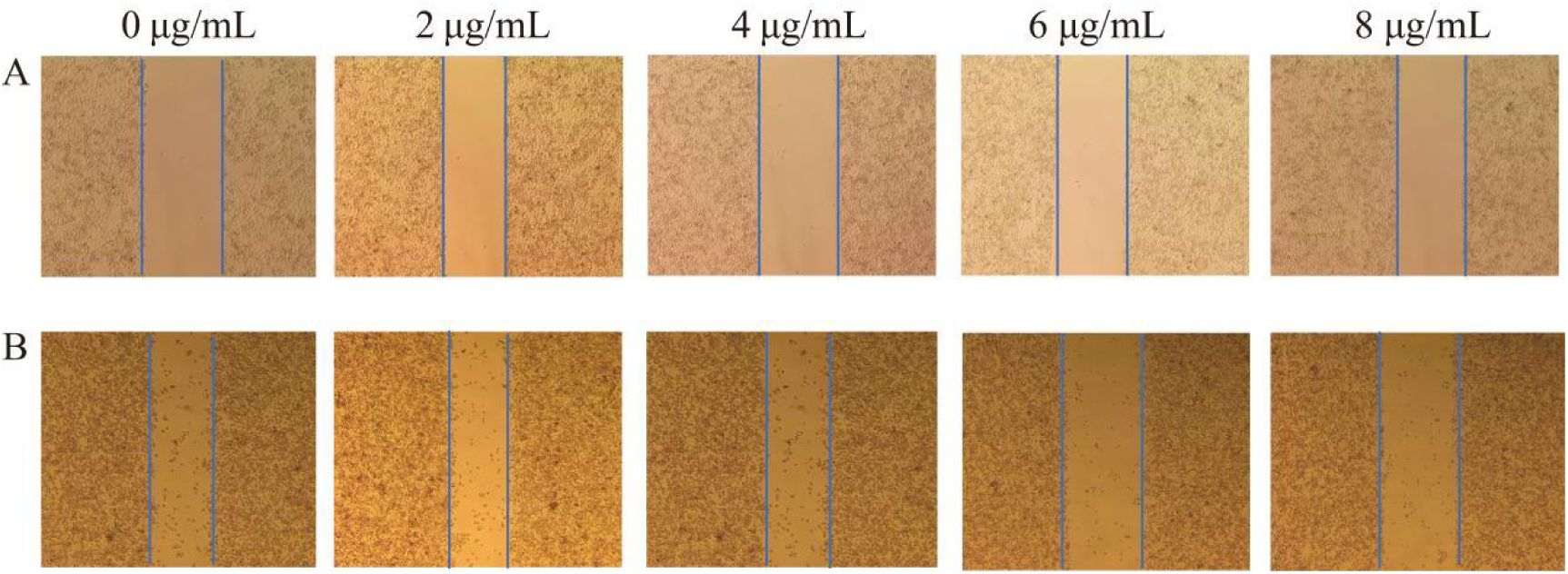
Melittin inhibits the migration of U14 cells *in vitro* at different concentrations (0, 2, 4, 6, and 8 μg/mL) (A) 0 h. (B) 16 hours post-treatment.

### 2.3 Assessment of cell apoptosis following melittin intervention

U14 cells treated with varying concentrations of melittin (0, 2, 4, 6, and 8 μg/mL) were analyzed utilizing Annexin V-FITC/PI double staining and flow cytometry. As illustrated in Figure 5, control cells (0 μg/mL) and those exposed to 2 μg/mL melittin exhibited high viability, with the vast majority of cells residing in the live population quadrant (Q1-LL). However, escalating concentrations of melittin induced a profound, dose-dependent shift in cell populations(**Figure 3A**). The flow cytometric scatter plots and corresponding quantitative analysis revealed that melittin predominantly triggered late apoptosis (Q1-UR quadrant, Annexin V+/PI+). Specifically, the proportion of late apoptotic cells increased to 10.59% at 4 μg/mL, and surged dramatically to 29.29% and 32.01% following exposure to 6 μg/mL and 8 μg/mL melittin, respectively. Throughout this process, the early apoptotic population (Q1-LR) remained relatively low across all concentrations. These flow cytometric data unequivocally demonstrate that melittin exerts its potent cytotoxic effect on U14 cells primarily by driving robust, dose-dependent late apoptosis(**Figure 3B**).

**Figure 3.**
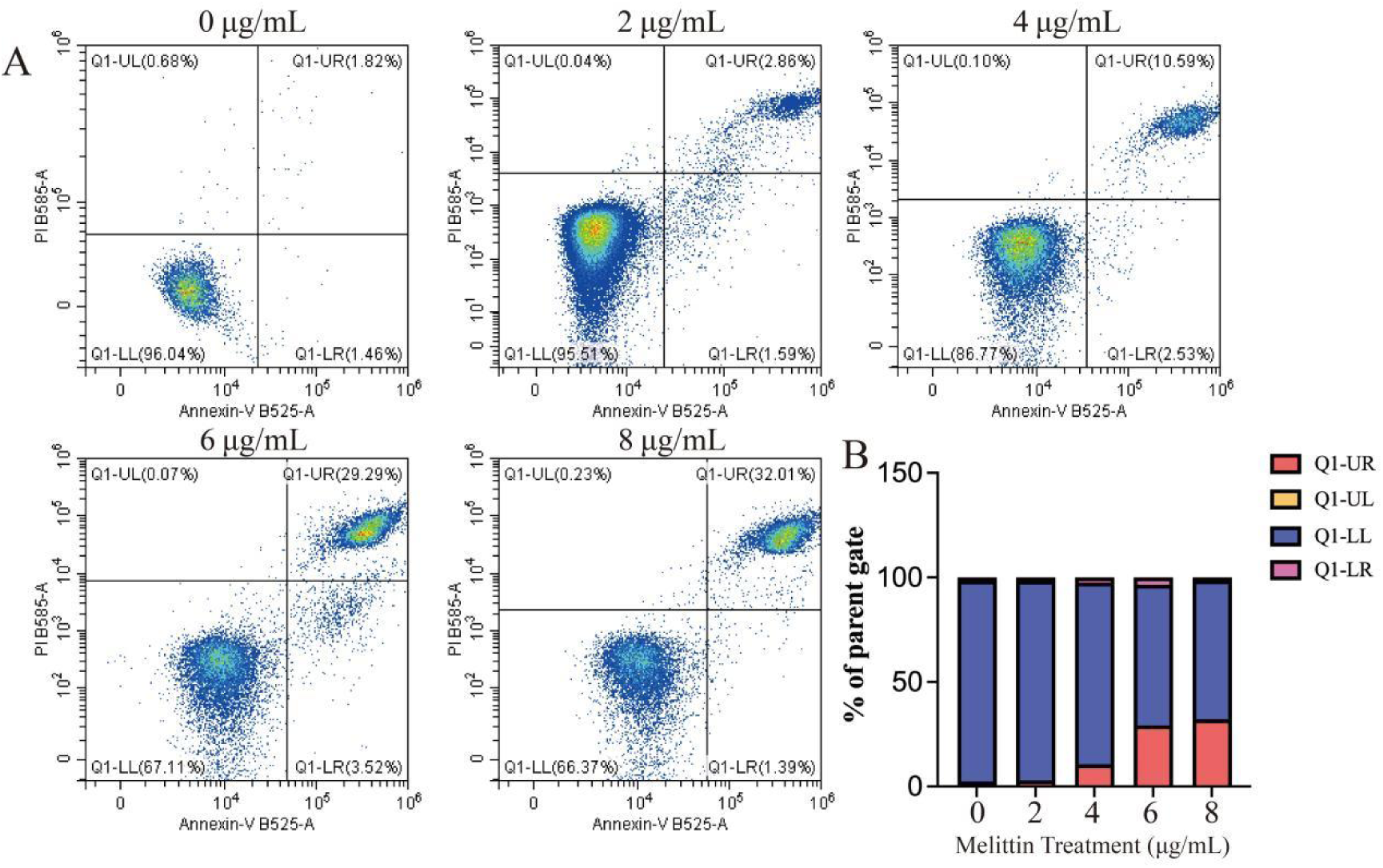
Melittin induces dose-dependent apoptosis in U14 cells. (A) Representative Annexin V-FITC/PI flow cytometry plots following melittin treatment (0, 2, 4, 6,8 μg/mL). Quadrants denote necrotic (UL), late apoptotic (UR), live (LL), and early apoptotic (LR) populations. (B) Quantitative percentages of the respective cell sub-populations.

**Figure 4.**
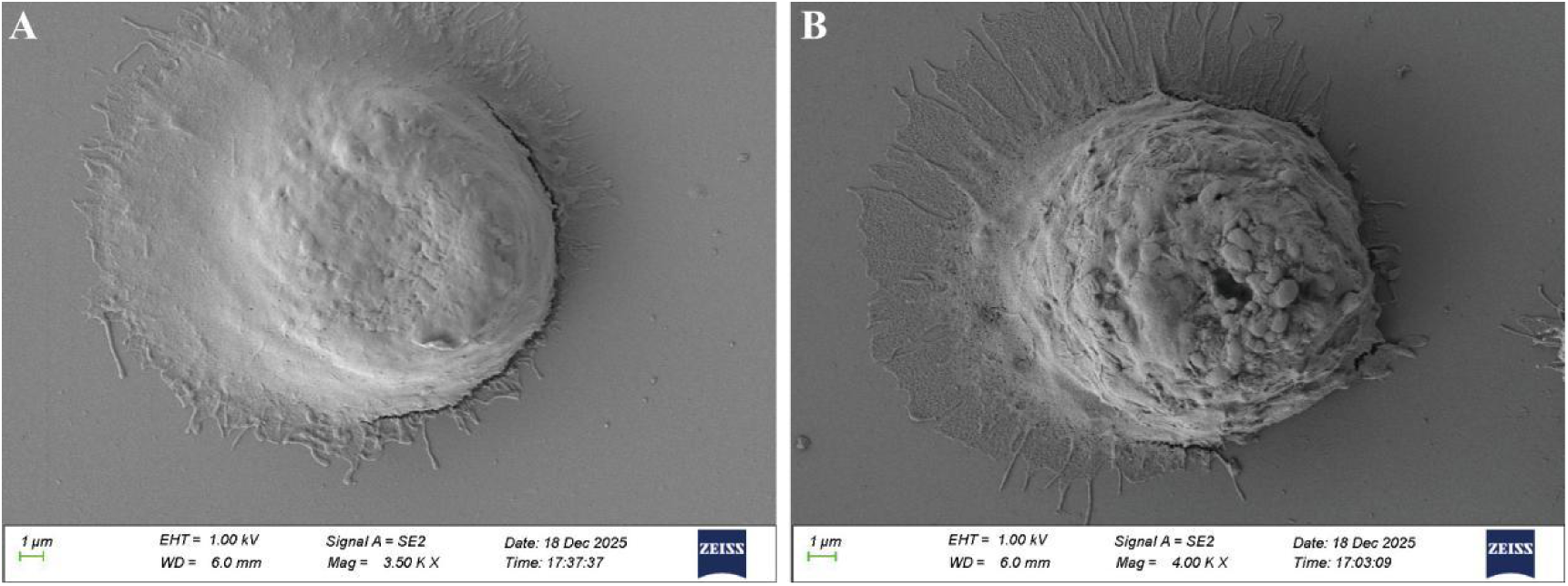
Detection of melittin-induced cell surface ultrastructural alterations in U14 cells under scanning electron microscopy. (A) Un-treated control group, (B) Melittin-treated group at 4 μg/mL.

**Figure 5.**
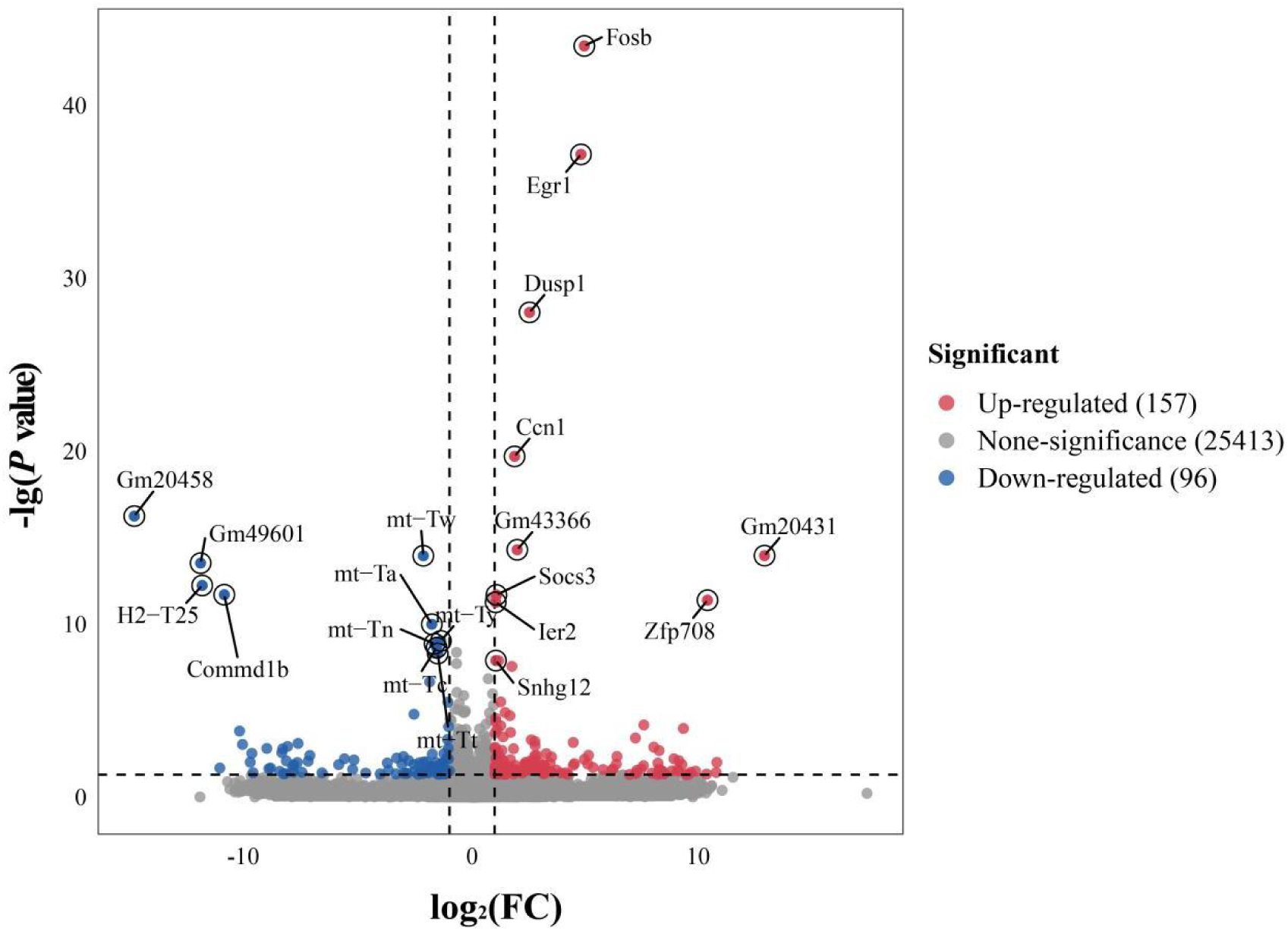
Volcano diagram of differentially expressed genes in murine U14 cells following melittin treatment

### 2.4 Observation of cellular morphology and ultrastructure following melittin intervention

By using high-resolution SEM, a distinct morphological contrast between the vehicle-treated and melittin-treated cells was onserved (**Figure 4**). Control cells (**Figure 4A**) exhibited an intact membrane architecture and normal surface topography. In contrast, following melittin exposure, the treated cells exhibited altered topographical features, characterized by prominent membrane blebbing, surface granulation, structural collapse, and extensive membrane perforations (**Figure 4B**).

### 2.5 Differential expression pattern of genes in melittin-treated U14 cells

Comparative analysis identified a total of 254 differentially expressed genes (DEGs) between the melittin-treated and control groups, consisting of 158 up-regulated and 96 down-regulated genes. Specifically, *Fos* (log_2_FC = 6.10, *Q* < 0.001), *Fosb* (log_2_FC = 4.96, *Q* < 0.001), and *Egr1* (log_2_FC = 4.81, *Q* < 0.001) emerged as the top three up-regulated genes. Conversely, the most pronounced down-regulation was observed in the uncharacterized genomic loci *Gm20458* (log_2_FC = −14.94) and *Gm49601* (log_2_FC = −12.01), as well as the mitochondrial tRNA gene *mt-Tw* (log_2_FC = −2.16, *Q* < 0.001) (**Figure 5**).

### 2.6 Functional and Pathway Enrichment Analyses of DEGs

GO analysis identified significant enrichment in biological process (BP) terms related to RNA polymerase II transcriptional regulation, signal transduction, differentiation, immunity, and apoptotic regulation (Figure 6, Table S1). Other enriched BP terms included GPCR signaling and responses to xenobiotics, IL-6, and IL-17. Cellular component (CC) mapping localized these DEGs to the membrane, nucleus, mitochondrion, endoplasmic reticulum, and Golgi apparatus, while molecular function (MF) was dominated by DNA-binding transcription factor activities (**Figure 6A**).

**Figure 6.**
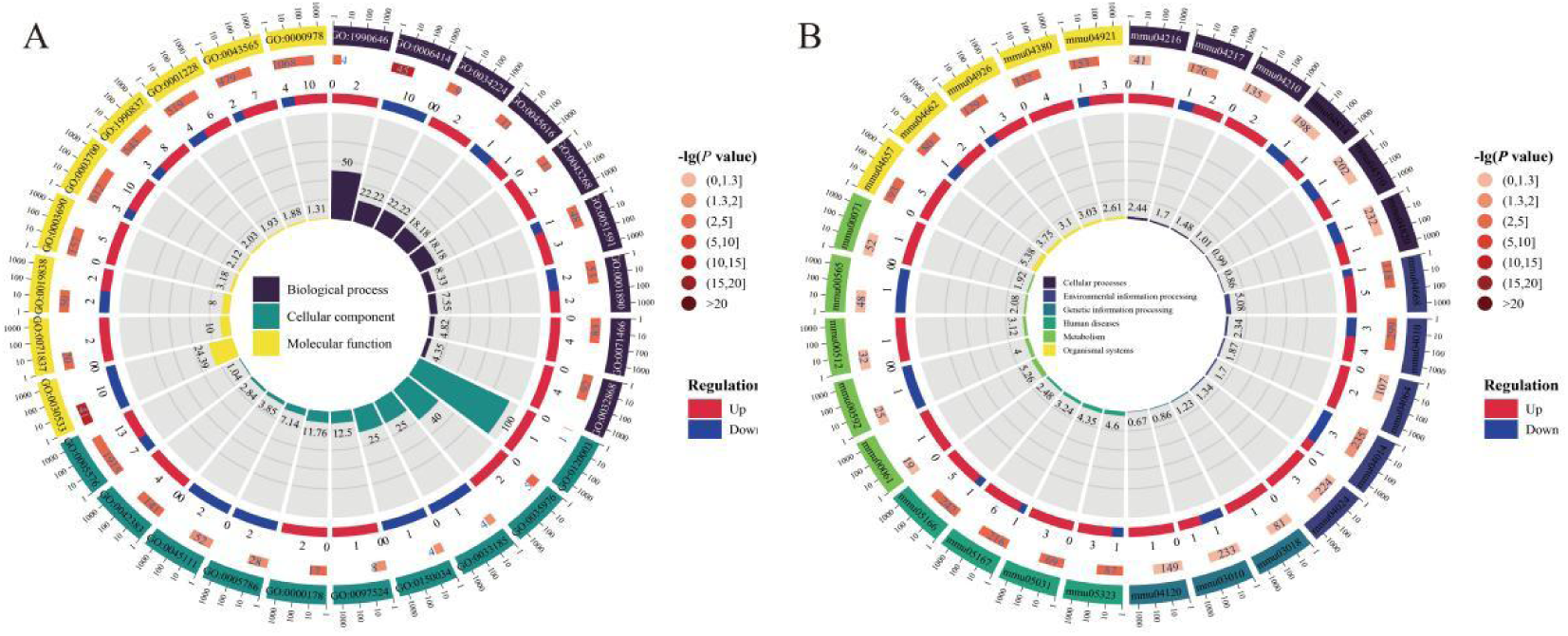
Bubble plots of GO terms(A) and KEGG pathways(B) enriched by differentially expressed genes in murine U14 cells following melittin treatment

KEGG profiling linked DEGs predominantly to signaling pathways involved in cellular stress responses and immune regulation, enriching in the TNF, IL-17, and MAPK pathways. Gene-level mapping highlighted the significant up-regulation of Immediate early genes (IEGs) (*Fos*, *Fosb*, *Jun*, and *Egr2*). *Fos* and *Fosb* were mapped to the enriched Activated Protein 1 (AP-1) complex, while *Jun* was involved in MAPK/TNF-driven proliferative and inflammatory cascades. *Egr2* was linked to sequence-specific DNA binding. Additionally, the ciliary-associated gene *Tulp2* was up-regulated, whereas the angiogenic regulator *Etv2* was markedly down-regulated (**Figure 6B**).

### 2.7 Reactome Pathway Enrichment Analysis

To systematically decode the transcriptomic alterations at the pathway level, Reactome enrichment analysis was conducted, yielding 20 significantly enriched biological cascades (Figure 7). The functional landscape revealed a pronounced divergence between up- and down-regulated gene modules. Notably, the Formation of AP-1 complex pathway displayed the most profound statistical significance (*P* = 0.0039, *Q* = 0.02, Rich factor ≈ 0.95), driven exclusively by up-regulated genes. Additional cascades dominated by up-regulated genes encompassed specific metabolic and receptor adaptations, such as Myoglobin binds oxygen and CHRM4 binds acetylcholine.

**Figure 7.**
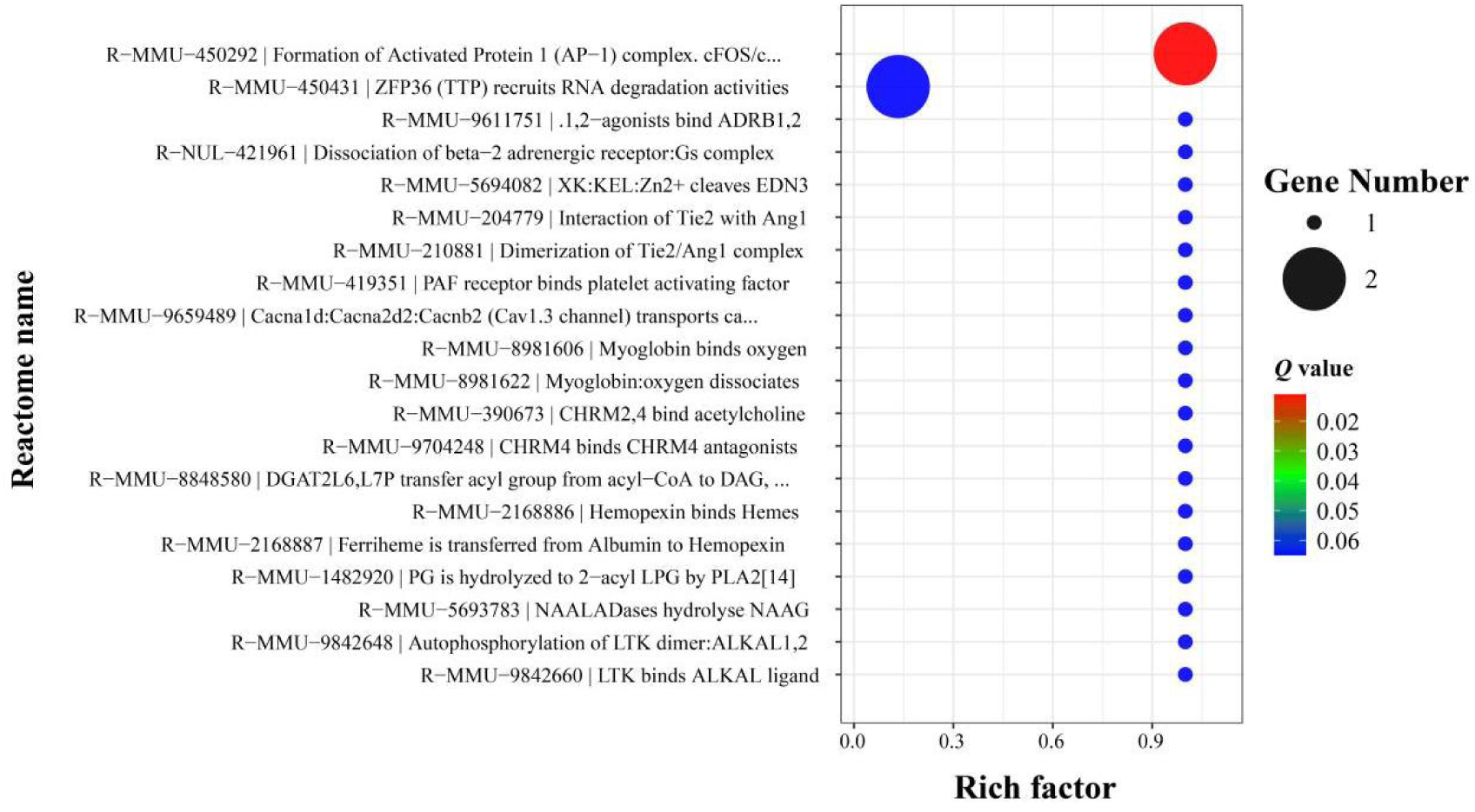
Bubble plot of reactome pathways enriched by differentially expressed genes in melittin-treated murine U14 cells

Conversely, a distinct functional module characterized by down-regulated genes was predominantly mapped to RNA turnover and angiogenesis-related signaling. This included significant enrichment of ZFP36 (TTP) recruits RNA degradation activities alongside classical angiopoietin-Tie2 signaling networks (e.g., Interaction of Tie2 with Ang1 and Dimerization of Tie2/Ang1 complex). Furthermore, cascades involving the autophosphorylation of LTK dimers, beta-adrenergic receptor signaling, and platelet-activating factor receptor signaling also reached statistical significance. Taken together, these data objectively delineate a transcriptomic signature defined by the induction of AP-1-associated networks and the concurrent suppression of Tie2-mediated signaling cascades (**Figure 7**).

### 2.8 GSEA of DEGs

Global transcriptomic profiling via GSEA demonstrated a significant suppression of translation-related gene sets following melittin treatment. The most negatively enriched GO terms were structural constituent of ribosome (NES = −3.16, FDR < 0.001) and translational elongation (NES = −3.15, FDR < 0.001), alongside triplet codon-amino acid adaptor activity (NES = −3.07, FDR < 0.001) and cytoplasmic translation (NES = −2.95, FDR < 0.001) (**Figure 8**).

**Figure. 8.**
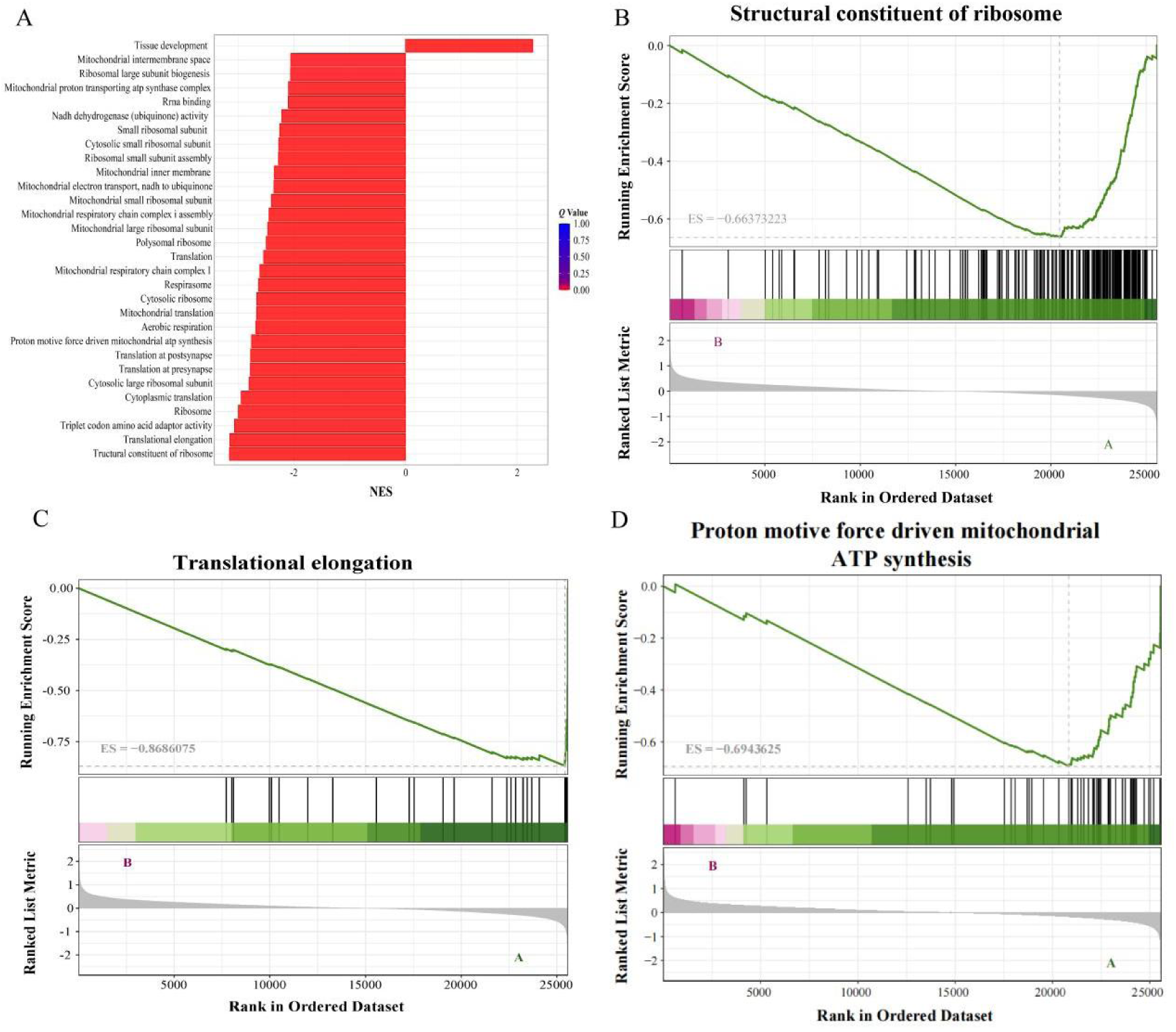
Gene set enrichment analysis of translation- and energy metabolism-related gene sets in cervical cancer cells following melittin treatment (A) Bar chart of negatively and significantly enriched GO terms (B-D) Representative enrichment plots of core down-regulated functional gene sets associated with structural constituent of ribosome, translational elongation, and proton motive force-driven mitochondrial ATP synthesis.

Consistent with the GO analysis, KEGG pathway GSEA identified Ribosome (NES = −3.34, FDR < 0.001). This was closely followed by a substantial reduction in the Oxidative phosphorylation pathway (NES = −2.42, FDR < 0.001). Furthermore, significant negative enrichment was observed in networks controlling proteasomal degradation and those associated with neurodegenerative diseases (NES from −1.62 to −1.88, FDR < 0.05) (**Figure 9**).

**Figure. 9.**
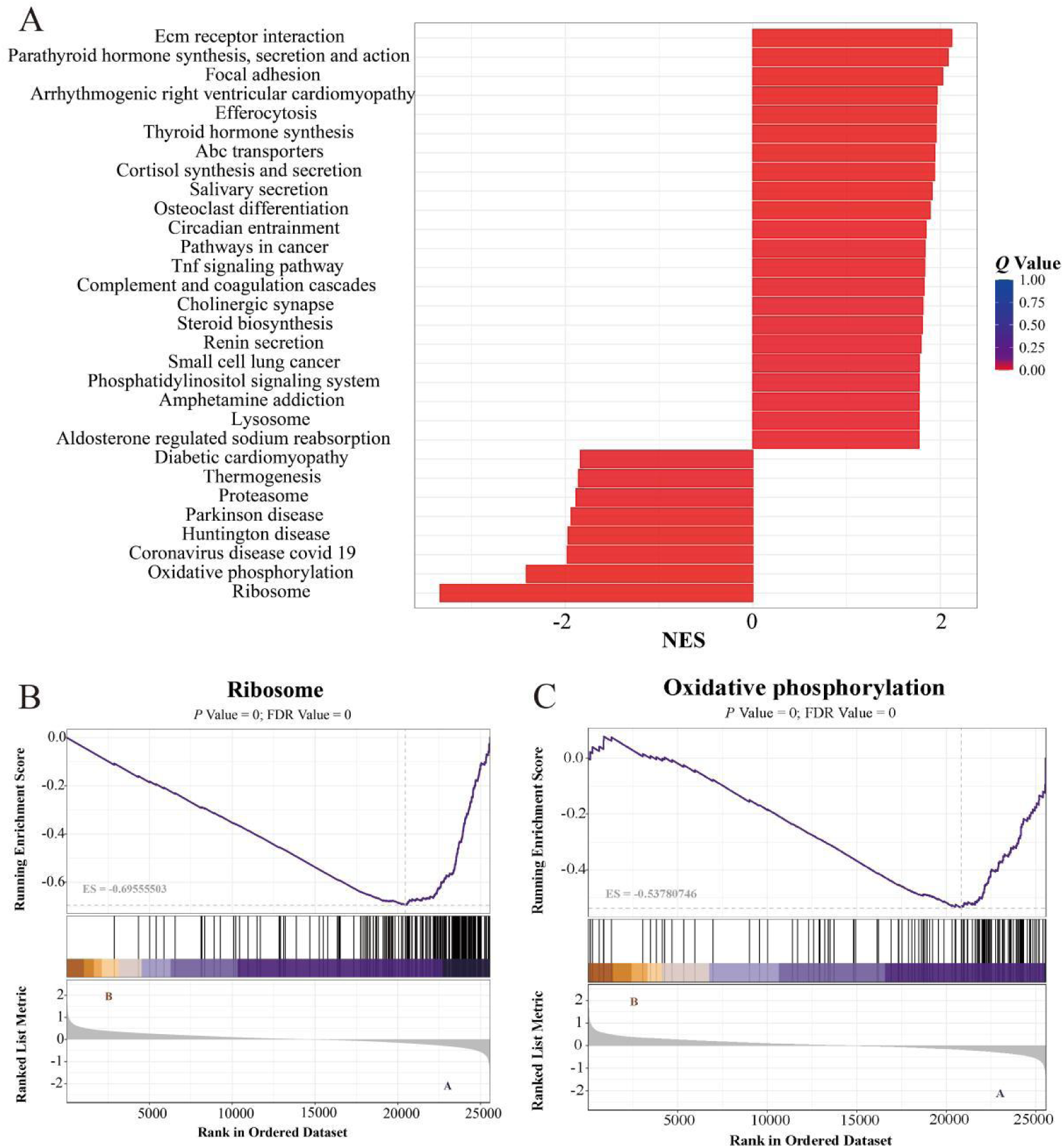
KEGG pathway-based GSEA uncovers repression of core translational and metabolic pathways in melittin-treated cervical cancer cells. (A) Horizontal bar plot of enriched KEGG pathways (B-C) Representative GSEA enrichment plots for the two core down-regulated pathways: ribosome pathway and oxidative phosphorylation pathway.

### 2.9 PPI Network Analysis of DEGs

Topological evaluation of the PPI network delineated a highly centralized and densely interconnected regulatory module. Within this network hierarchy, Jun emerged as the principal hub node, exhibiting the highest degree of connectivity (degree = 22). Furthermore, a distinct sub-network of immediate interacting partners, predominantly consisting of *Fos*, *Fosb*, *Egr1*, *Dusp1*, *Ccn1*, and *Zfp36*, densely aggregated directly around this central *Jun* anchor (**Figure 10**).

**Figure. 10.**
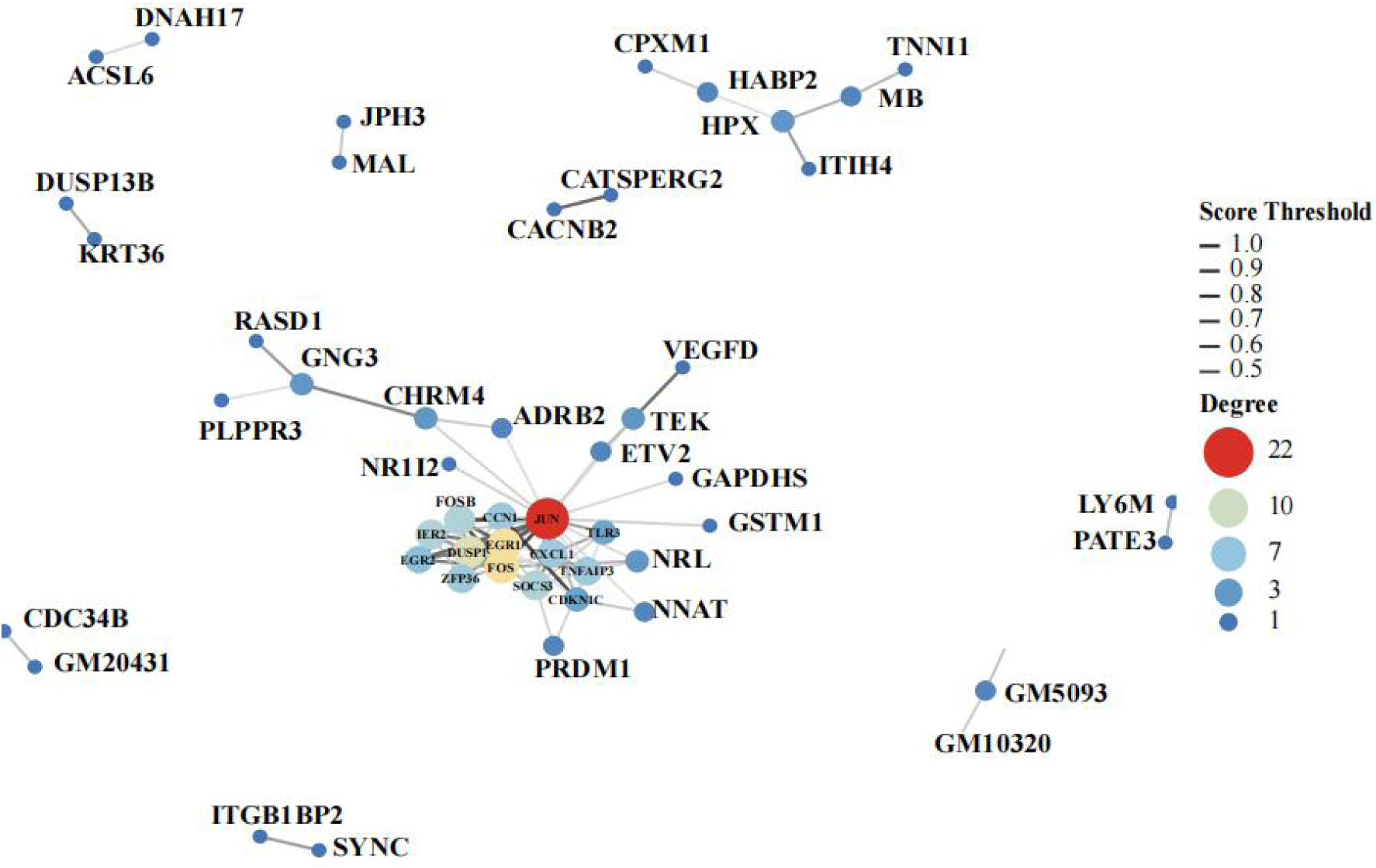
Protein-protein interaction network of differentially expressed genes in melittin-treated murine U14 cells.

### 2.10 RT-qPCR Validation of Transcriptome data

RT-qPCR assay showed a profound, highly significant transcriptional burst of *Fos* (*P* < 0.0001), accompanied by a significant concurrent induction of *Fosb* (*P* < 0.05). Concurrently, although *Egr*, *Jun*, and *Prdm1* displayed a trend toward up-regulation and *Nr1i2* trended down-regulation, these alterations did not reach statistical significance. The results verified the reliability of transcriptome data used in this study (**Figure 11**).

**Figure 11.**
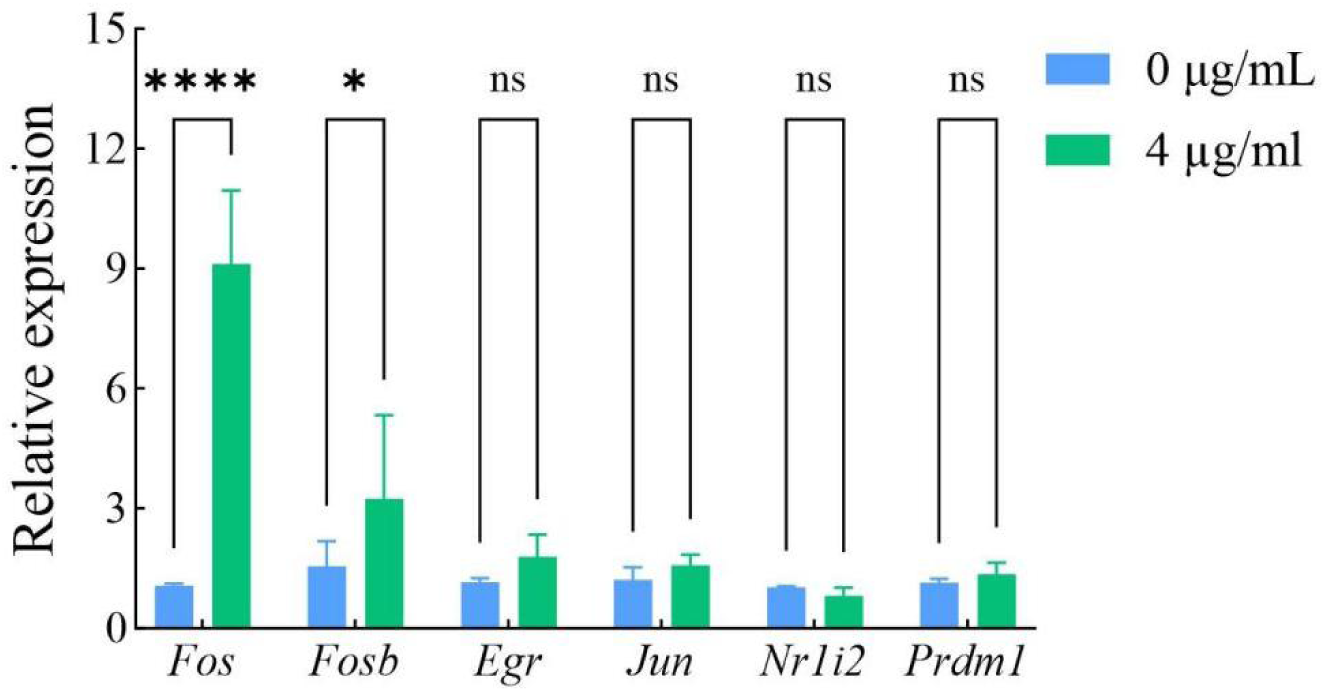
Relative mRNA expression levels of six target genes under 4 μg/mL treatment. Data are presented as mean ± SD. Different lowercase letters above the bars indicate significant differences (^ns^*P* > 0.05; \**P* < 0.05; \*\*\*\**P* < 0.0001 (Two-way ANOVA).

## 3 Discussion

Cervical cancer represents one of the most prevalent gynecological malignancies worldwide, and the clinical limitations of conventional chemoradiotherapy highlight the urgent need for novel, highly effective therapeutic agents ^[17]^. In recent years, natural antimicrobial peptides have garnered significant attention in oncology. Melittin, the principal active component of honeybee venom, has emerged as a particularly potent candidate due to its renowned membrane-disrupting properties and broad-spectrum anti-tumor activities ^[18–19]^. Despite its well-documented cytotoxicity across various solid tumors, the profound intracellular molecular perturbations and the global transcriptomic reprogramming induced by melittin in cervical cancer remain inadequately elucidated ^[9,20–21]^. To address this knowledge gap, the present study comprehensively investigated the anti-cancer efficacy of melittin in murine U14 cervical cancer cells by integrating macroscopic phenotypic characterizations with in-depth transcriptomic profiling.

Phenotypically, melittin exhibited potent, dose-dependent cytotoxicity against U14 cells, profoundly suppressing both acute viability and long-term clonogenic survival (**Figure 1**). Flow cytometry confirmed that this lethal effect was primarily driven by the rapid induction of late apoptosis (**Figure 3**). Ultrastructural analysis via SEM provided direct visual evidence of this process, revealing severe membrane blebbing, structural collapse, and extensive perforation. These physical disruptions are consistent with melittin’s amphipathic nature, which facilitates its rapid insertion into lipid bilayers to execute membrane lysis ^[22,23]^ (**Figure 4**). Furthermore, higher concentrations of melittin virtually abolished cell motility in wound scratch assays. This robust anti-migratory effect aligns with recent pharmacological findings across various solid tumors ^[24,25,26]^, confirming melittin’s capacity to restrict cervical cancer progression (**Figure 2**).

To elucidate the underlying molecular alterations, GO enrichment analysis provided initial insights into the global transcriptomic shift. The predominant enrichment in biological processes governing RNA polymerase II-mediated transcription, signal transduction, and apoptotic regulation indicates that the physical disturbance caused by melittin rapidly translates into a systemic nuclear transcriptional response (**Figure 6A**). Following this, KEGG pathway topological profiling highlighted significant enrichment in environmental information processing networks, notably the TNF, IL-17, and MAPK signaling pathways. Gene-level mapping within these pathways revealed a striking up-regulation of IEGs, including *Fos*, *Fosb*, *Jun*, and *Egr2*. While MAPK and TNF signaling cascades are traditionally associated with cell proliferation under physiological conditions (**Figure 6B**), recent transcriptomic insights reveal that the acute hyperactivation of IEGs under severe external intervention often signifies an unresolvable cellular stress response that drives apoptosis rather than survival ^[27,28]^. Concurrently, the marked down-regulation of *Etv2*, a key regulator enriched in angiogenesis, suggests that melittin simultaneously suppresses pro-angiogenic signals crucial for tumor progression.

To further dissect these transcriptomic shifts, Reactome enrichment analysis was employed, uncovering a distinct divergence in pathway functionality. On one hand, melittin overwhelmingly triggered the formation of AP-1 complex pathway, firmly establishing the AP-1 transcription factor complex as the primary regulatory hub governing the cellular stress response. On the other hand, it precipitated a significant decline in modules related to RNA degradation and the angiopoietin-Tie2 signaling cascade. The Tie2/Ang1 axis is widely recognized as a fundamental driver of endothelial recruitment, vascular maturation, and microenvironmental remodeling in cervical cancer ^[29–30]^. Therefore, the profound transcriptomic suppression of this network perfectly corroborates the anti-migratory phenotype observed in our cellular assays, pointing to a severe disruption in essential tumor-stroma crosstalk(**Figure 7**).

To overcome the threshold-dependent biases of conventional DEG profiling, we utilized GSEA to capture system-wide functional shifts. GSEA revealed a sweeping, highly significant suppression of translation- and energy metabolism-related gene sets. Pathways governing ribosomal biogenesis and mitochondrial oxidative phosphorylation exhibited profound negative enrichment. Rapidly proliferating cancer cells maintain an absolute biosynthetic and energetic dependency on hyperactive translation and oxidative ATP production ^[31–32]^ (**Figure 8-9**). Emerging cancer therapeutic strategies heavily emphasize targeting these exact vulnerabilities ^[33]^. The global collapse of ribosomal architecture and oxidative phosphorylation indicates that melittin severely impairs the translational and energetic capacities of U14 cells, depriving them of the metabolic pillars required for survival.

Further, topological evaluation of the PPI network delineated a highly centralized regulatory interactome, structurally reinforcing our functional enrichment findings. Anchored by Jun as the primary topological hub, this network features a dense sub-cluster containing *Fos*, *Fosb*, *Egr1*, *Dusp1*, *Ccn1*, and *Zfp36* (**Figure 10**). Crucially, our subsequent RT-qPCR quantitative validation further refined this regulatory landscape. While global RNA-seq identified a broad engagement of the early response network, targeted RT-qPCR analysis uncovered that this acute stress response is fundamentally driven by a profound and highly significant transcriptional burst specific to *Fos* and *Fosb* (**Figure 11**). In contrast, alterations in other network nodes (such as *Jun*, *Egr*, and *Prdm1*) exhibited non-significant trends, underscoring *Fos* and *Fosb* as the principal responsive targets under this specific melittin treatment regimen. The concentrated hyperactivation of this *Fos*-dominated AP-1 axis, acting in concert with key stress regulators like *Dusp1* and *Zfp36* ^[34–35]^, orchestrates an overwhelming cellular stress response that dictates irreversible apoptosis.

In summary, melittin effectively eradicates murine U14 cervical cancer cells through a highly synchronized, multi-targeted mechanism. The robust anti-tumor efficacy is driven by direct plasma membrane lysis, followed by a sweeping transcriptomic collapse of ribosomal biogenesis and oxidative metabolism, Tie2-mediated migratory arrest, and fatal AP-1 stress network hyperactivation. Collectively, these molecular insights highlight melittin as a superior therapeutic candidate for cervical carcinoma.

## Conclusion

In summary, this study systematically validates the anti-cervical cancer efficacy of melittin through a dual-level investigation. Experimentally, melittin exerts potent cytotoxicity by directly disrupting the plasma membrane and effectively abolishing cell migration and survival. Mechanistically, downstream transcriptomic profiling and RT-qPCR validation reveal that this physical damage initiates a massive intracellular crisis, highlighted by the profound suppression of ribosomal translation and oxidative metabolism, the blockade of Tie2-mediated pro-angiogenic signals, and the lethal induction of a *Fos/Fosb*-dominated AP-1 stress interactome. Ultimately, this multi-modal mechanism, combining structural, metabolic, and transcriptional targeting—provides a robust scientific rationale for developing melittin as a clinical therapeutic agent against cervical carcinoma.

## Appendix

**Table S1.** GO term and KEGG pathway enrichment analyses of the differentially expressed genes in melittin-treated U14 cells.

| Category | Term/Pathway | *P* value | *Q* value | Nmuber |
| --- | --- | --- | --- | --- |
| BP | Regulation of transcription by RNA polymerase II | 0.004 | 0.089 | 16 |
| BP | Cellular response to xenobiotic stimulus | 0.001 | 0.089 | 4 |
| BP | Cellular response to interleukin-17 | 0.003 | 0.089 | 2 |
| BP | Positive regulation of interleukin-6 production | 0.003 | 0.089 | 4 |
| BP | Cell differentiation | 0.029 | 0.147 | 10 |
| BP | Immune system process | 0.316 | 0.417 | 5 |
| BP | Apoptotic process | 0.455 | 0.537 | 4 |
| BP | G protein-coupled receptor signaling pathway | 0.992 | 0.996 | 5 |
| CC | Membrane | 0.515 | 0.584 | 49 |
| CC | Nucleus | 0.918 | 0.934 | 28, |
| CC | Mitochondrion | 0.078 | 0.205 | 15 |
| CC | Endoplasmic reticulum | 0.120 | 0.252 | 13 |
| CC | Golgi apparatus | 0.618 | 0.672 | 7 |
| MF | DNA-binding transcription factor activity | < 0.001 | 0.014 | 13 |
| KEGG | TNF signaling pathway | <0.001 | 0.002 | 6 |
| KEGG | IL-17 signaling pathway | <0.001 | 0.003 | 5 |
| KEGG | MAPK signaling pathway | <0.001 | 0.013 | 7 |

